# QTL mapping and proteomic profiling of barley: insights into resistance and susceptibility to *Pyrenophora teres* f. *teres*

**DOI:** 10.64898/2026.02.04.703889

**Authors:** Buddhika A. Dahanayaka, Richard Wilson, Sadegh Balotf, James Hane, Anke Martin

**Author notes:** Corresponding authors &.

## Abstract

*Pyrenophora teres* f. *teres (Ptt),* the causal agent of net form net blotch disease in barley, is an economically important fungal pathogen worldwide. Understanding both host resistance mechanisms and pathogen virulence factors is essential for developing durable net form net blotch resistant barley cultivars. Quantitative trait loci (QTL) mapping was conducted using a cross between two *Ptt* isolates, one virulent on the barley cultivar ‘Prior’ and the other being avirulent. A major QTL associated with virulence on Prior was detected on chromosome 5. A progeny isolate possessing this QTL, together with the two parental isolates, was subsequently used in the proteomic analyses. Label-free proteomics was used to quantify *in planta* the protein profile changes in Prior following inoculations with the virulent and avirulent parental *Ptt* isolates, and the virulent progeny isolate. Leaf samples were collected at two (D2) and five (D5) days post-inoculation, and proteomic analyses performed to identify proteins associated with host resistance and pathogen virulence. A dataset comprising 2,886 barley proteins and 51 *Ptt* proteins was analysed. Principal component analysis (PCA) of the barley Prior proteomes revealed distinct clustering based on resistance and susceptibility at D5, while D2 samples formed a separate cluster. The PCA of the *Ptt* proteomes identified separate clusters, one comprised of the D2 and D5 avirulent parental isolate and another cluster of the virulent isolates at D5 only. Gene ontology analysis of the Prior proteins that were significantly increased in the resistant compared to the susceptible groups revealed functional categories related to protein translation, biosynthesis and chloroplast activities. The proteins that were significantly increased in the susceptible compared to the resistant Prior group were associated with organic acid and carbohydrate metabolism. The proteomic profiles and bioinformatic analysis generated in our study provide novel insights into the molecular basis of resistance and virulence in the barley-*P. teres* pathosystem.

**Key message:** This study reveals the first *in planta* proteomic profiles of both barley and *Pyrenophora teres* f*. teres*, identifying unique virulence-associated proteins and host responses linked to resistance and susceptibility.

## Introduction

Net blotch is one of the most economically significant diseases affecting barley (*Hordeum vulgare*) globally (Smedegard-Petersen 1976). Yield losses due to net blotch can reach up to 40% in susceptible barley cultivars (Murray&Brennan 2010). The fungal species *Pyrenophora teres* [anamorph *Drechslera* (Sacc.) Shoemaker] is the causal agent of net blotch in barley, which exists as two forms, *P. teres* f. *teres (Ptt*) causing net form net blotch (NFNB) and *P. teres* f. *maculata* (*Ptm*) causing spot form net blotch (SFNB). *Pyrenophora teres* f. *teres* and *P. teres* f. *maculata* cause net-like reticulate and spot-like circular necrotic lesions on barley leaves, respectively. The two forms of *P. teres* are morphologically similar but genetically distinct (Ellwood et al. 2012), hence molecular markers are required to distinguish the two forms from each other (Poudel et al. 2017). Population genetics and pathotyping studies of *Ptt* have revealed significant genetic diversity among global populations, with numerous distinct pathotypes contributing to the complexity of the barley–*Ptt* pathosystem (Fowler et al. 2017; Ellwood et al. 2019; Dahanayaka et al. 2021b; Bakonyi et al. 2024).

*Pyrenophora teres* f*. teres* is a necrotrophic pathogen, with an initial short biotrophic phase followed by a predominant necrotrophic phase (Lightfoot&Able 2010). Pathogens like *Ptt* may employ both biotrophic (nutrition from live host cells) and necrotrophic (nutrition from dead host cells) infection strategies to invade the host successfully. Virulence-related proteins secreted by pathogens can act as pathogen-associated molecular patterns (PAMPs), which are detected by host-encoded pattern recognition receptors (PRRs). This recognition initiates PAMP-triggered immunity (PTI), representing the first layer of innate plant defence (De Wit et al. 1997; Jones&Dangl 2006; De Wit et al. 2009).

Studies show that fungal pathogens produce effectors that supress PTI (Friesen et al. 2007; Leiva-Mora et al. 2024; Thynne et al. 2024). These effectors enable the pathogen to invade the host plant, activate effector-triggered immunity (ETI) and induce disease reactions in the host (De Wit et al. 2009). Effector recognition by host plants initiates a co-evolutionary dynamic often described as an “arms race,” where pathogens adapt through random mutations and natural selection, leading to effector loss or modification, or the evolution of new effectors that suppress host’s ETI (De Wit et al. 1997; Jones&Dangl 2006; De Wit et al. 2009). In necrotrophic pathogens, effector-host interactions often follow an inverse gene-for-gene model, where effector recognition promotes susceptibility or compatible interaction with the host. These interactions are governed by both qualitative and quantitative inheritance, indicating a more complex model than the classical gene-for-gene concept (Friesen et al. 2008). Multiple approaches such as QTL mapping to identify resistance loci in the host or virulence loci in the pathogen, proteomics analyses to characterise host/pathogen proteins and RNA sequencing to profile transcriptional responses during infection to identify host resistance and pathogen virulence genes have been used to elucidate the barley-*Ptt* pathosystem. However, the pathosystem remains complex due to polygenic resistance, diverse pathogen effectors and as a result molecular basis of the interaction between *Ptt* and barley remains unclear (Clare et al. 2020).

Proteomics has been used to study the virulence of several major plant-pathogen species, including: *Parastagonospora nodorum* (Casey et al. 2010), *Magnaporthe oryzae* (Meng et al. 2019; Wang et al. 2024), *Leptosphaeria maculans* (Marra et al. 2010), *Fusarium graminearum* (Paper et al. 2007; Rauwane et al. 2020)*, Botrytis cinerea* (González-Fernández et al. 2014), *Sclerotinia sclerotiorum* (Li et al. 2018; Tian et al. 2021), *Ustilago* maydis (Böhmer et al. 2007; Weiland&Altegoer 2021), and *Rhizoctonia solani* (Anderson et al. 2016). Pathogen-focussed proteomic studies tend to improve gene annotation and/or infer potential virulence functions, whereas host-focussed studies compare broad responses to infection between resistant and susceptible cultivars. To date, only one previous study has used proteomics to investigate the *Ptt*-barley pathosystem *in planta*, in which the apoplast proteins from healthy and infected barley leaves were compared (Hassett et al. 2020).

In the current study, label-free shotgun proteomics was used to investigate the *in planta* molecular interaction between *Ptt* and barley cultivar “Prior”. The primary goal was to characterize both host and pathogen protein profiles during infection and to compare temporal changes following inoculation with virulent and avirulent *Ptt* isolates. By identifying key proteins and pathways involved with virulent/susceptible (compatible) and avirulent/resistant (incompatible) interactions, this research enhances our understanding of barley defence mechanisms and *Ptt* virulence strategies. These insights could inform breeding programs aimed at improving durable net form met blotch resistance and to support the development of more effective disease management strategies.

## Materials and methods

### 1. Plant materials

Four barley genotypes, Beecher, Commander, Prior, and Skiff were used for phenotyping in a QTL mapping study. Beecher and Skiff served as avirulent controls and Commander as the virulent control.

Barley cultivar Prior was used for all *in planta* proteomic analyses conducted in this study. Seedling assays were conducted across a panel of 24 barley cultivars/genotypes (Prior, Beecher, Skiff, Maritime, Algerian, Ciho 11458, Ciho 5791, Corvette, Gilbert, Harbin, Skipper, Kombar, Tallon, Yerong, Harrington, Fleet, Buloke, Commander, Hindmarsh, Vlamingh, Navigator, Shepherd, Rosalind and Fathom) to determine the virulence phenotypes of the three *Ptt* isolates and to understand the effect of chromosome 5 QTL on different barley genotypes. Seed of all barley cultivars/genotypes used in this study were provided by the Grain Research and Development Cooperation (GRDC), Australia.

### 2. Fungal materials

For QTL mapping analysis, a bi-parental mapping population (P2) of 88 individuals was developed by crossing *Ptt* isolates *P451* with *P155*. Isolates *P451* and *P155* are progeny from a cross (P1) between *Ptt* isolates NB81 and HRS09127 used in the *Ptt* MP-NAM study by Dahanayaka &Martin (2023). According to the Dahanayaka &Martin (2023) study, isolate *P451* was virulent on Prior and possessed a major QTL on chromosome 5 associated with Prior virulence while *P155* was avirulent on Prior with no associated QTL. Population P2 was developed to examine the segregation of the major virulence QTL on chromosome 5 and to obtain single-QTL isolates that possessed only this major QTL. The use of single-QTL isolates enabled the identification of corresponding host target proteins associated with resistance and susceptibility, minimising confounding effects from minor QTL.

For protein analysis, we used the parental isolates of P1 (*P451* and *P155*), and a progeny isolate from P2, *SP23*.

#### Fungal crossing and ascospore collection for fungal mapping population

The fungal cross was established according to the method described by Dahanayaka et al. (2021a). Five 50 mm long autoclaved pieces of barley straw were placed onto Sach’s agar (Hebert 1971) plates before the agar had set. An approximately 25 mm^2^ mycelial plug was excised from cultures of *P451* and *P155* grown on half-strength potato dextrose agar (PDA; 20g/L PDA; Biolab Merck, Darmstadt, Germany) plates and placed adjacent to the barley straw. The crossing plates were kept in transparent plastic bags to prevent desiccation and placed into an incubator at 15°C with a 12h light/12h dark photoperiod.

Once ascomata had matured, lids of Petri plates were replaced with 2% water agar plates for ascospore collection. Plates were monitored daily for ascospores. Single ascospores were collected from water agar plates under a dissecting microscope (SMZ 745; Nikon, Melville, New York) and transferred to PDA plates using a sterile glass needle. Water agar plates were replaced with new plates each day.

#### Genotyping and QTL mapping

Fungal isolates were grown on half-strength PDA media at 22°C for 10 days. Mycelium from each of the fungal isolates was gently scraped off the plate and freeze dried for 24h in a 2mL tube. Freeze dried samples were sent to Diversity Arrays Technology Pty. Ltd. (Canberra, ACT, Australia) for DNA extraction and DArTseq™.

The genetic linkage map was constructed as described previously (Dahanayaka et al. (2022a); Dahanayaka &Martin (2023). SilicoDArT and SNP marker data obtained from DArTseq™ were filtered for non-polymorphic marker and those having more than 10% missing data. Using MapManager QTXb20 version 2.0 (Manly et al. 2001) markers were grouped into linkage groups with a *p* = 0.05 search linkage criterion. RECORD (Van Os et al. 2005) was used to order the markers. The final genetic map was obtained after manually curating the genetic groups (Lehmensiek et al. 2009). Marker sequences (62 bp) were aligned with the reference genome W1-1 (BioSample SAMEA4560035 available under PRJEB18107 BioProject) using the bowtie2 (Langmead&Salzberg 2012) function in Galaxy to compare marker positions.

QTL analyses were conducted using the composite interval mapping method in Windows QTL Cartographer version 2.5 (Wang 2012). For each trait, experiment-wise logarithm of the odds (LOD) threshold values at the 0.05 significance level were determined through 1000 permutation tests (Churchill&Doerge 1994; Doerge&Churchill 1996).

#### Phenotyping of the P. teres f. teres isolates

Phenotypic assessment of *Ptt* progeny was conducted in a controlled environment room (CER) at the University of Southern Queensland (UniSQ), Toowoomba, Australia, with three replicates as described in Dahanayaka et al. (2022a). Barley genotypes were grown for 14 days with 12h day and night cycles at 23°C and at 17°C, respectively.

Inoculum was produced according to the method described for *P. tritici-repentis* (Jacques et al. 2021), briefly, where one-day old mycelial plugs on V8 medium (150mLV8® vegetable juice, 15g agar, 3g CaCO_3_, 800mL distilled water)were exposed to near-UV light for 24h to induce sporulation, followed by 24h dark incubation to promote conidia maturation. Spores were harvested in a Tween 20 solution to produce hypha-free suspensions. Conidial concentrations were determined using a haemocytometer and adjusted to a final concentration of 10,000 conidia/mL.

After 14 days each pot consisting of four barley genotypes was inoculated with 2.5mL of conidial suspension. Inoculated plants were incubated in the dark for 24h at 95% humidity with a temperature of 23 ± 1°C and were transferred back to the CER for nine days. Disease reaction scores on the second leaf were scored nine days after inoculation (Tekauz 1985) on a 1 (highly resistant) to 10 (highly susceptible) scale.

For virulence phenotyping of the single QTL *SP23* progeny isolate and the parental isolates 24 barley cultivars listed under “Plant materials” were grown in pots with 5cm diameter and 14cm height and the phenotyping was performed as mentioned in the above section using the same environment conditions.

### 3. Proteomic analysis

#### Sample preparation for proteomic analysis

Inoculation of the barley cultivar Prior was conducted following the completely randomized block design in a CER at the UniSQ, Australia (Dahanayaka et al. 2022b). Sixteen seeds of Prior were grown in each pot with 5cm diameter and 14cm height at 20 ± 1°C and day and 16 ± 1°C night for 14 days.

The conidial suspension of each isolate was prepared for plant inoculation as described in Dahanayaka et al. (2022b). Once the second leaves of the barley plants were fully expanded, i.e. fourteen days after planting, 3mL of conidial suspension (10,000 conidia/mL) were sprayed onto the plants in each pot. Inoculated pots were incubated in the dark for 24h at 95% humidity with a temperature of 23 ± 1°C. After 24h, plants were transferred to the same controlled environment room mentioned above. The second leaves from the inoculated plants were collected at 2 and 5 days post-inoculation (dpi). Three replicates (three leaves each from three pots) of each sample were frozen in liquid nitrogen immediately after harvest and stored in - 80°C until protein extraction.

Proteins were extracted from three independent replicates of each isolate using 50-70mg of frozen leaf samples and adding 650µL extraction buffer [7M urea, 2M thiourea, 1% dithiothreitol (DTT), 100mM NaCl, 40mM Tris, pH8.0]. The samples were homogenised for 30s using a Fast Prep-24 bead beater (MP Biomedicals, Seven Hills, NSW, Australia) at 5,500rpm at room temperature and then centrifuged at 16,000rpm for 10 min at 4°C. The supernatant was transferred to a 1.5mL tube. Six volumes of cold acetone were added to the resulting clarified solution and samples were incubated overnight at −20°C. The samples were centrifuged at 10,000rpm for 8min at 4°C. The resultant protein pellets were left to air dry for 5min at room temperature before resuspending them in the same extraction buffer mentioned above. Protein concentration was determined using the Pierce™ 660 nm protein assay (Thermo Fisher Scientific Pty Ltd, VIC, Australia). A total of 30µg of protein from each sample was then used for downstream analyses mentioned below.

Samples were reduced overnight at 4°C using 10mM DDT followed by alkylation using 50mM iodoacetamide at room temperature in the dark for 2h. The Single-Pot Solid-Phase-enhanced Sample Preparation (SP3) method (Hughes et al. 2019) was used for the trypsin/LysC digestion of the proteins. The resulting peptides were then desalted using ZipTips (Merck, Darmstadt, Germany).

#### Peptide analysis by liquid chromatography and mass spectrometry

The desalted peptides were dried in a SpeedVac concentrator (miVac Duo, GeneVac Australia and dissolved in 12µL 0.05% (w/w) trifluoroacetic acid in water/acetonitrile (98:2). Approximately μg of peptide per sample was analysed using an Ultimate 3000 nano RSLC system coupled to an Orbitrap Q-Exactive HF mass spectrometer (Thermo Scientific, Waltham, MA, USA). Samples were separated over 2h LC gradients and analysed using data-independent mass spectrometry (DIA-MS) according to the parameters described previously (Balotf et al. 2021). Briefly, peptides were separated using PepMap 100 C18 preconcentration and analytical columns (75μm x 2cm and 25cm, respectively, Thermo Scientific, Waltham, MA, USA) using a sample trapping/elution configuration. Samples were first loaded onto the preconcentration column at 5μL/min prior to separation at 300nL/min. DIA-MS parameters were: 2.0kV spray voltage, S-lens RF level of 60 and heated capillary set to 250°C. MS1 spectra (390–1240 m/z) were acquired in profile mode at a resolution of 120,000 (AGC target set to 3 x 10^6^), followed by 26 DIA x 25amu MS2 scans over the range of 397.5–1027.5m/z, using 1amu overlap between sequential windows. MS2 spectra were acquired at a resolution of 30,000 in centroid mode, using an AGC target of 1 x 10^6^, a maximum injection time of 55ms and normalized collision energy set to 27. Raw DIA-MS files were imported into Spectronaut software (v 17, Biognosys) and searched against a combined FASTA file composed of the UniProt *Ptt,* 0-1 (Ellwood et al. 2010) and barley, Morex (Mayer et al. 2012) protein sequence databases (UP000001067 and UP000011116, respectively) accessed on March 3^rd^, 2023, and comprising 11,705 and 34,528 entries, respectively. Essentially, default Biognosys (BGS) settings were used for both library generation and extraction of MS1 and MS2 DIA profiles. Database search parameters included oxidized methionine and N-terminal acetylation as variable modifications and carbamidomethyl of cysteine as a fixed modification. Relative protein abundance across samples was achieved by targeted DIA analysis of the resulting spectral library, including MS2-level peptide quantitation, protein quantitation based on the maxLFQ algorithm and cross-run global normalization based on median peptide intensities.

#### Statistical analysis – barley proteomic data

The protein intensity data were separated into two datasets based on the species assigned to the proteins: *P. teres* and Prior (barley) proteins. The Prior protein dataset was filtered by removing proteins with more than one missing value per sample group. Normalized protein intensity values were transformed (log_2_) followed by imputing the missing values with a normal distribution using Perseus v2.0.9.0 (Tyanova et al. 2016), according to default settings.

Differentially abundant proteins were identified using analysis of variance (ANOVA) to compare multiple treatment groups or Student’s *t*-test for pairwise comparison using an FDR (false discovery rate) threshold of 0.05 and s0 setting of 0.1. Principal component analysis (PCA) plots were used to visualise the relationship between treatment groups, based on the subset of ANOVA-significant proteins, using Perseus (v2.0.9.0).

Proteins that showed significant changes in abundance were subjected to gene ontology (GO) enrichment analysis using ShinyGO (Ge et al. 2020) which assigns proteins to functional categories and biological pathways based on curated databases. STRING database (https://string-db.org/) based on a threshold of FDR < 0.05 was used to construct association networks, allowing to infer pathway-level enrichment from statistically overrepresented GO terms among the differentially abundant proteins.

#### Statistical analysis – Pyrenophora teres f. teres proteomic data

The *Ptt* protein dataset was filtered by removing proteins with two or more missing values in each sample group, but proteins identified in the basis of a single peptide match were retained. This filtered set of *Ptt* proteins was then manually screened to identify proteins that were uniquely present in all three replicates of the virulent parental and virulent progeny compared to avirulent parental isolates, to identify proteins that may be important for the virulence of *P. teres*. For statistical analysis, normalized protein intensity values were transformed (log_2_) followed by imputation of missing values prior to PCA and representation of the data as a heat map using hierarchical cluster analysis using Perseus (v2.0.9.0). Pairwise comparison between sample groups was used to identify proteins which were significantly (Student’s *t*-test FDR < 0.05, s0 = 0.1) altered in abundance. Functional enrichment of differentially abundant proteins was performed using the STRING database (https://string-db.org/) based on a threshold of FDR < 0.05. Gene ontology analyses were conducted using ShinyGO (Ge et al. 2020). EffectorP 3.0 (Sperschneider&Dodds 2022) was used to predict potential effector proteins which may be important for the infection process of *P. teres*.

## Results

### Genetic map and QTL mapping

The genetic map of population P2 consisted of 12 linkage groups containing 316 markers in total with linkage groups having between 3 and 64 markers (Table 1) (Supplementary_QTL mapping). The targeted linkage group, chromosome 5 contained 38 markers with an average distance of 5.5 cM between the flanking markers. Disease reaction scores of the progeny phenotyped on Prior ranged from 1 to 8.5 (Figure 1A) (Supplementary_QTL mapping). Disease reaction scores of the parental isolates *P155* and *P451* were 2.5 and 7.5, respectively and *SP23* had a score of 8. QTL analysis detected a QTL on chromosome 5 for Prior virulence with a LOD score of 16 and a phenotypic variation explained of 75% (Figure 1B). Parent *P451* contributed the virulence. Isolate *SP23* was virulent on Prior and possessed the QTL on chromosome 5. This isolate plus the parents *P451* and *P155* were used for the proteomic analysis.

**Table 1.** Genetic map information of the *Pyrenophora teres* f*. teres P451/P155* cross.

| Chromosome | Number of markers | Size cM | Average distance between flanking markers (cM) |
| --- | --- | --- | --- |
| 1 | 46 | 132.0 | 2.9 |
| 2 | 21 | 87.7 | 4.2 |
| 3 | 8 | 19.9 | 2.5 |
| 4 | 26 | 52.5 | 2.0 |
| 5 | 38 | 209.1 | 5.5 |
| 6 | 41 | 116.5 | 2.8 |
| 8 | 33 | 79.4 | 2.4 |
| 9 | 64 | 50.8 | 0.8 |
| 10 | 3 | 11.9 | 4.0 |
| 11 | 14 | 64.6 | 4.6 |
| 12 | 3 | 3.4 | 1.1 |
| Unlinked | 64 |  |  |

**Figure 1.**
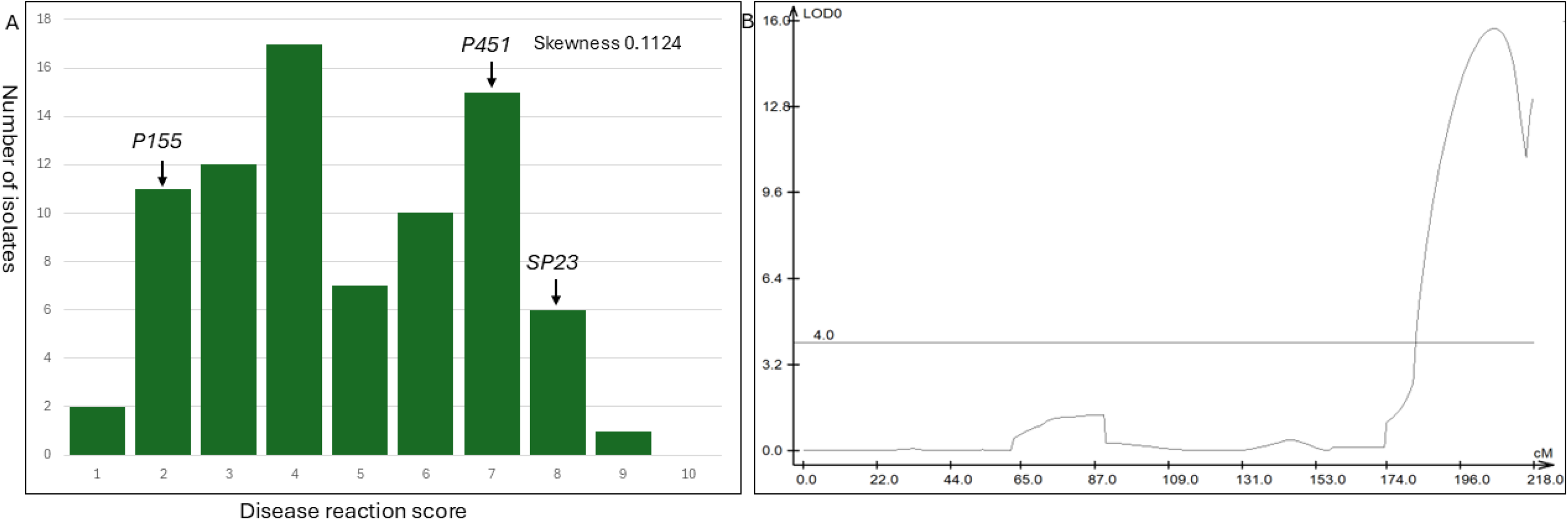
Disease reaction scores of *Pyrenophora teres* f. *teres* isolates of the P2 population on barley cultivar Prior (A) Quantitative Trait Loci (QTL) mapping of *Pyrenophora teres* f. *teres* virulence in population P2 (B). The graph illustrates the significant QTL on chromosome 5 associated with virulence on barley genotype Prior. The threshold LOD value (LOD 4) is indicated by the grey horizontal line. Linkage distances are marked along the x-axis in centi Morgan (cM).

### Phenotyping of the P. teres f. teres isolates

Average disease reaction scores of the three isolates on a differential set of 24 barley genotypes ranged from 1 to 9.8 (Figures 2 and 3). The virulence profile of the progeny isolate *SP2*3 was similar to the parental virulent isolate *P451* except for Corvette and Kombar. For Corvette, although both parents were virulent (>6), the progeny isolate was avirulent (<6), whereas for Kombar, the opposite was observed. All isolates were virulent on Gilbert, Skipper, Tallon, Harrington, Commander, Hindmarsh, Navigator, Shepherd and Fathom.

**Figure 2.**
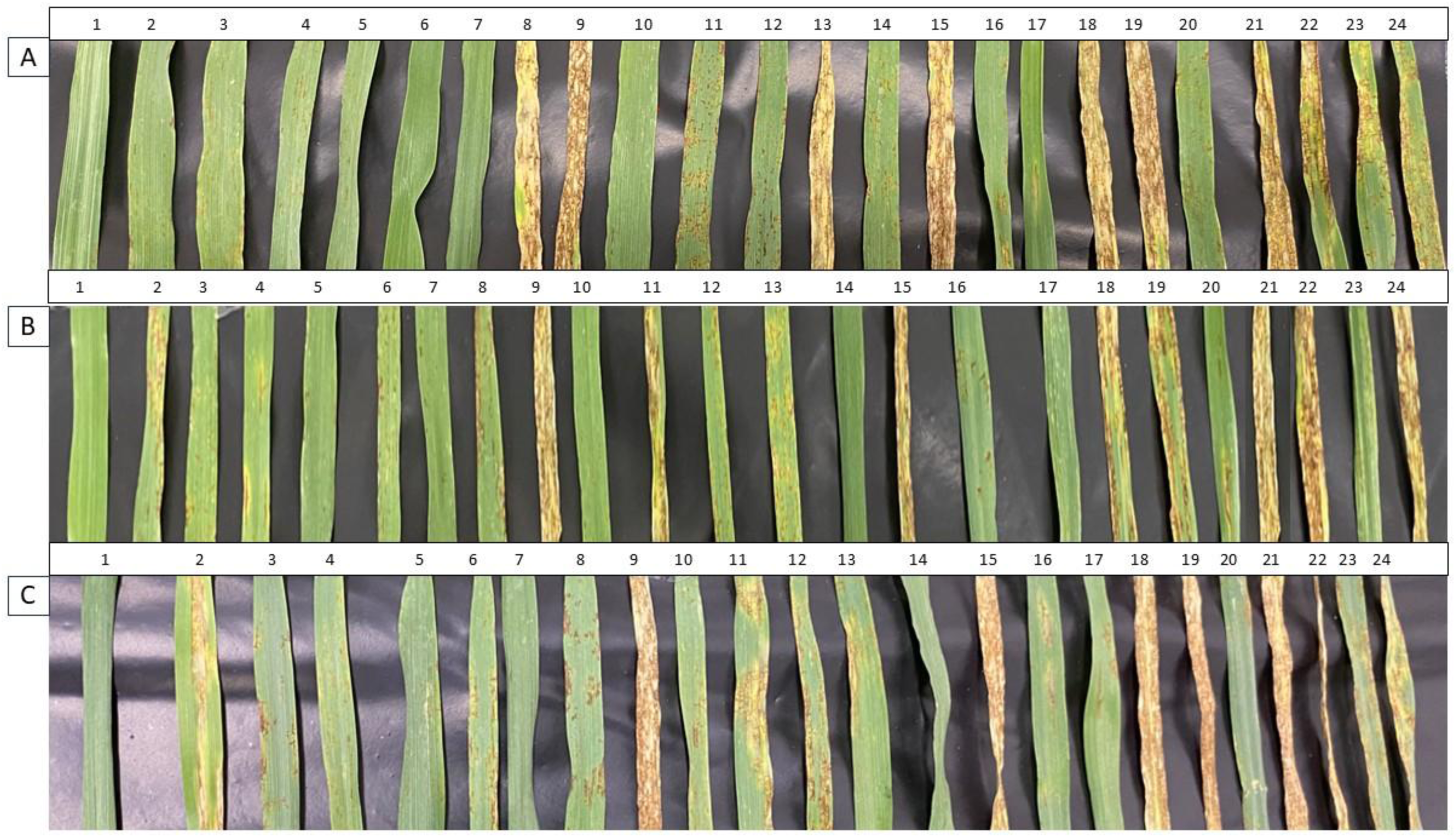
Disease symptoms caused by three *P. teres* f. *teres* isolates A. avirulent (*P155*) B. virulent (*P451*) and C. progeny (*SP23*) on 24 barley genotypes (1. Beecher, 2. Prior, 3. Skiff, 4. Maritime, 5. Algerian, 6. Ciho 11458, 7. Ciho 5791, 8. Corvette, 9. Gilbert, 10. Harbin, 11. Skipper, 12. Kombar, 13. Tallon, 14. Yerong, 15. Harrington, 16. Fleet, 17. Buloke, 18. Commander, 19. Hindmarsh, 20. Vlamingh, 21. Navigator, 22. Shepherd, 23. Rosalind and 24. Fathom).

**Figure 3.**
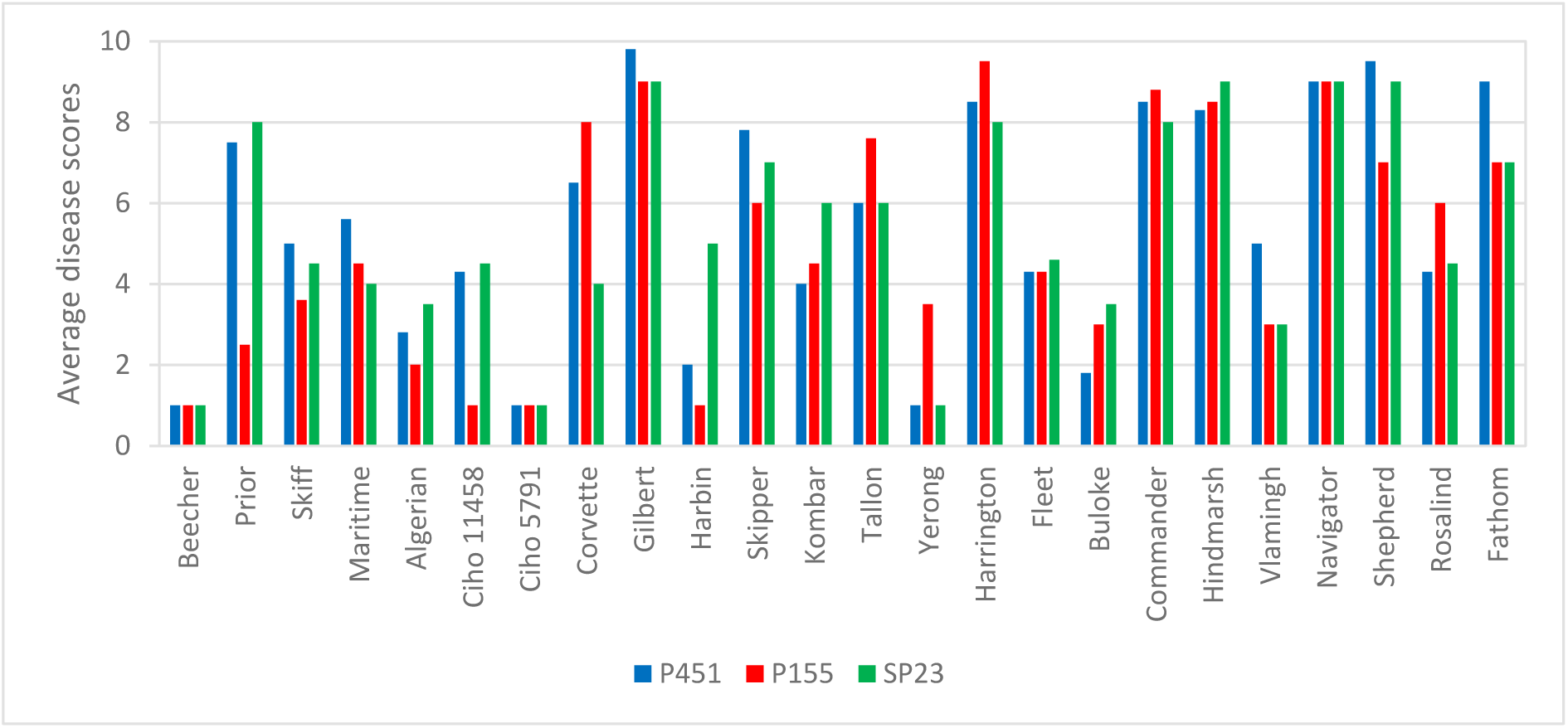
Disease reaction scores of *P. teres* f. *teres* isolates, avirulent (*P155*) virulent (*P451*) and progeny (*SP23*) on a differential set of 24 barley genotypes.

### The protein profiles of barley during P. teres f. teres infection

Proteomic analysis was performed on barley cultivar ‘Prior’ to identify host resistance-associated proteins in samples from incompatible interactions (host resistance to avirulent isolate *P155*) and pathogen virulence-associated proteins in samples from compatible interactions (host susceptibility to virulent isolates *P451* and *SP23*) at two (D2) and five days (D5) post-inoculation. Of the 3,502 proteins identified in total, 3,450 were of barley origin while 52 were unique to *P. teres* (Supplementary Table 1). After quality filtering (see Methods) 2,886 barley proteins were retained for analyses (Supplementary Table 2). Of these proteins, all but 13 were detected across all six treatments, therefore our comparison between groups was based on relative quantitation of mean protein intensity values rather than presence/absence of proteins between the different sample groups.

As multiple experimental conditions were performed in this study, we first used one-way ANOVA to test the effects of *Ptt* isolate virulence and inoculation time on the barley proteome. Using the subset of significantly different proteins (FDR < 0.05), principal component analysis (PCA) was performed to visualise relationships between sample groups. This was followed by post-hoc *t*-tests to compare sample groups exhibiting the greatest variation across the six treatments. Of the 2,886 barley proteins, 1,258 were differentially abundant across the D2 and D5 Prior samples inoculated with the *Ptt* isolates *P451, SP23* or *P155*, as determined by one-way ANOVA (FDR < 0.05). This set of proteins is provided in Supplementary Table 3.

According to PCA, the samples separated into three distinct clusters (Figure 4). One cluster comprised barley samples inoculated with all three *Ptt* isolates and collected at day 2 post-inoculation (cluster “D2”). This cluster clearly separated from the barley samples collected at day 5 which separated into virulent *Ptt* (*P451* and *SP23*; cluster “Susceptible_D5”) and avirulent *Ptt* (*P155*; cluster “Resistant_D5”) clusters. These three clusters were used to guide post-hoc *t*-test comparisons between groups.

**Figure 4.**
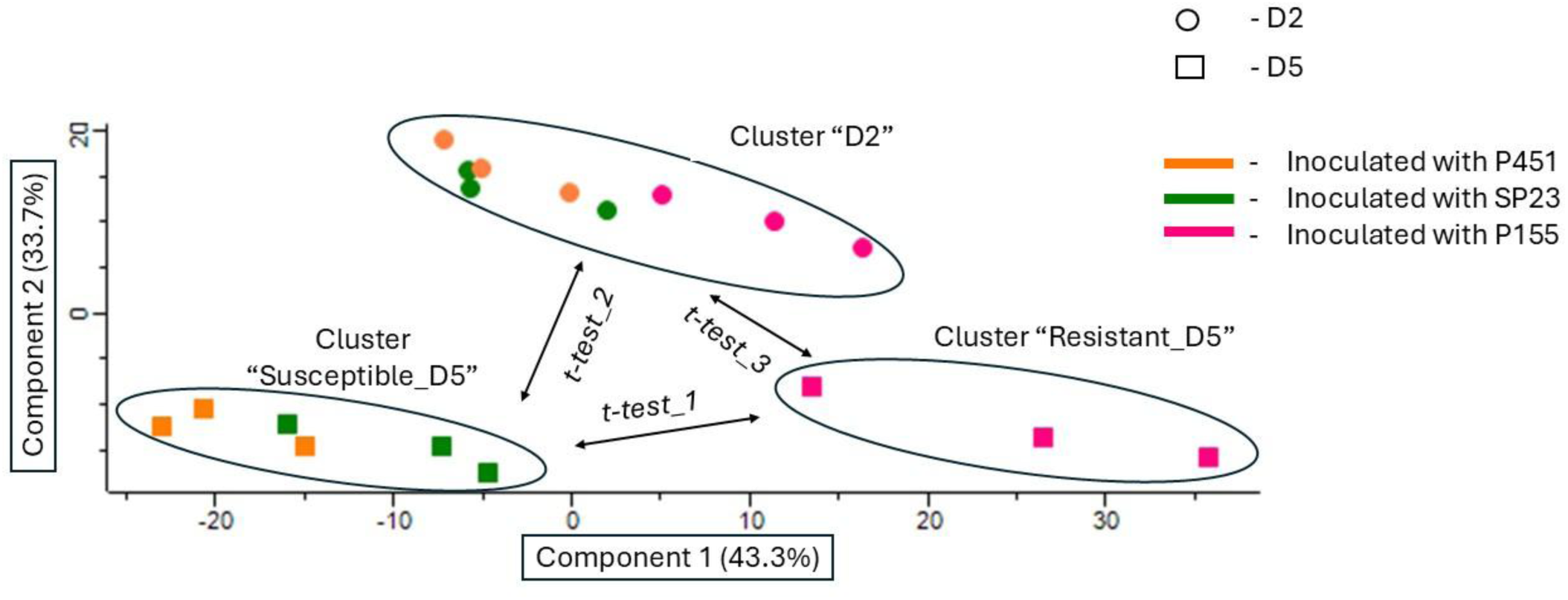
Principal component analysis plot based on the subset of proteins (n = 1,258) that were significantly altered in abundance (ANOVA) in the barley cultivar Prior proteome samples. The variations in the proteins of Prior inoculated with a virulent (*P451*) and an avirulent (*P155*) parental isolates of *Pyrenophora teres* f. *teres* (*Ptt*), as well as with a virulent progeny isolate (*SP23*) are shown. Clusters are labelled as Cluster “D2” (Prior proteome samples two days after inoculation), Cluster “Susceptible” (Prior proteome samples showed susceptible interaction with *Ptt*) and Cluster “Resistant” (Prior proteome samples showed resistant interaction with *Ptt*).

To identify a protein signature associated with overall *Ptt* susceptibility *vs* resistance, we compared the two sample groups comprising barley inoculated with virulent and avirulent *Ptt* isolates at D5 (clusters “Susceptible_D5” and “Resistant_D5” in Figure 4, respectively, *t*-*test_1*). The results of this *t-test* comparison are displayed as a volcano plot (Figure 5A) and the subset of 721 proteins showing significant differences in abundance is provided in Supplementary Table 4. Of these 721 proteins, 262 proteins were associated with *Ptt* resistance while 459 were associated with *Ptt* susceptibility. Proteins associated with *Ptt* resistance were important for various steps involved in plant defence mechanism such as cell signalling (calcium-dependent protein kinase), fungal cell wall catalytic enzymes (chitinase), lignan biosynthesis (dirigent), chloroplast metabolism (ferredoxins) and cellular detoxification (peroxidases and glutathione) (Supplementary Table 4).

**Figure 5.**
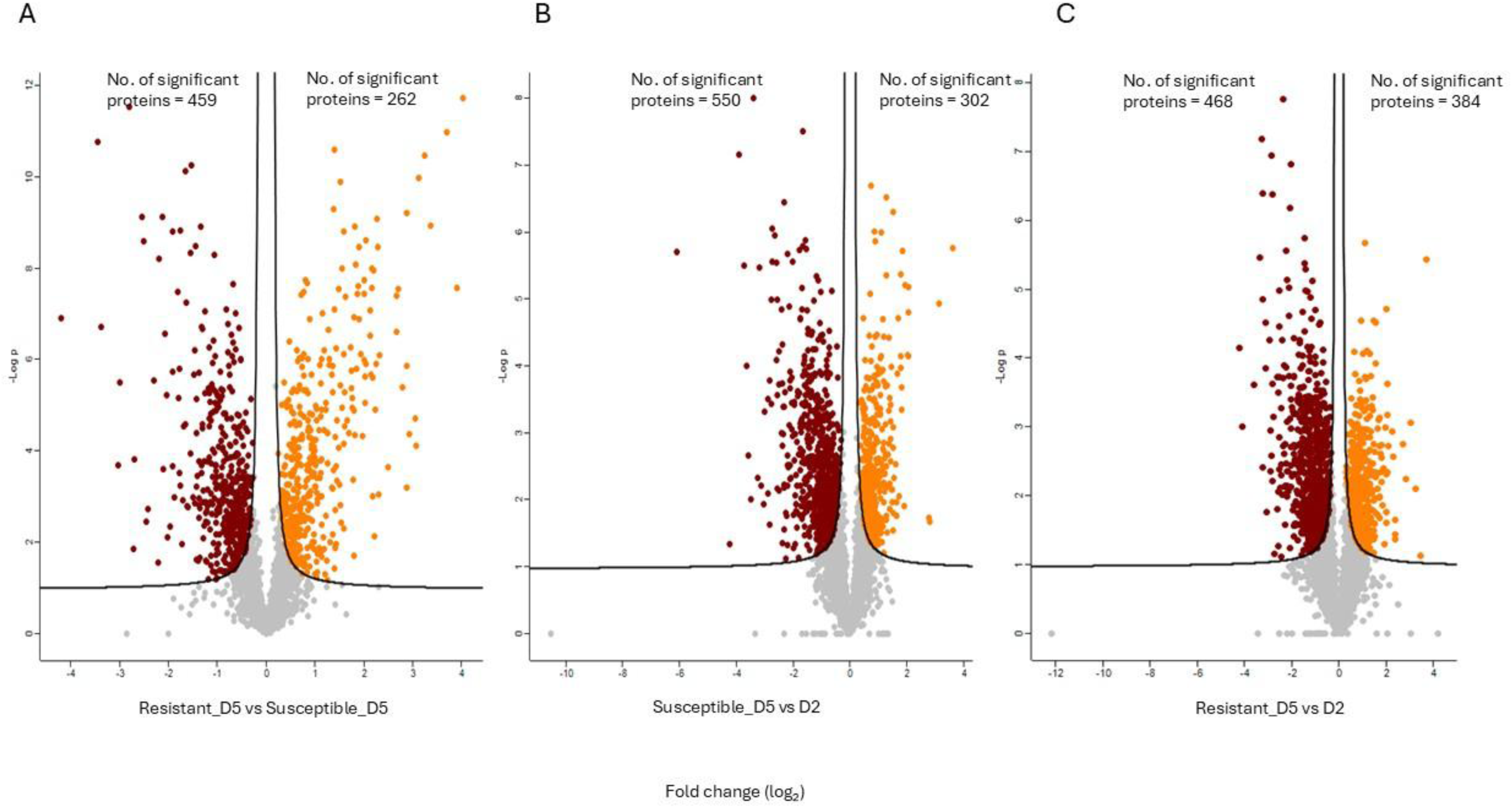
Volcano plots showing proteins with significant abundance changes between D5 and D2 in Prior proteomes inoculated with *P155* (avirulent *Ptt*), *SP23*, and *P451* (virulent *Ptt*). (A) Resistant D5 *vs* Susceptible D5; (B) Susceptible D5 *vs* D2; (C) Resistant D5 *vs* D2. Each plot displays log₂ fold change (x-axis) versus –log₁₀ p-value (y-axis). Proteins with significant changes (FDR < 0.05; s0 = 0.1) are highlighted: orange indicates increased abundance proteins, red indicates decreased abundance proteins, and grey represents non-significant proteins.

The PCA plot indicated a clear separation of barley proteomes at D2 and D5 post inoculation with *Ptt* (Figure 4). Therefore, to identify protein signatures that could represent initial (D2) and late (D5) responses of Prior to *Ptt* inoculation, we separately compared the samples in cluster D2 with samples inoculated with virulent *Ptt* (*P451* and *SP23*; cluster “Susceptible_D5”) and with the samples inoculated with avirulent *Ptt* (*P155*; cluster “Resistant_D5”) The results of these *t*-tests are displayed as volcano plots in Figure 5B (*t*-*test*_*2*) and Figure 5C (*t*-*test*_*3*), respectively. In both comparisons, subsets of 852 proteins were identified as significantly changed between the groups (Supplementary Tables 5 and S6). Several proteins identified in the barley proteome were classified as pathogenesis-related (PR) proteins belonging to different classes (Supplementary Tables 4, 5 and 6) (Ferreira et al. 2007; Gorjanović 2009). Notably, PR-2 proteins (Glucan endo-1,3-beta-D-glucosidases) exhibited increased abundance in Susceptible_D5 samples compared to Resistant_D5 and D2. PR-3, PR-4, PR-8, and PR-11 classes (chitinases) showed increased abundance in Resistant_D5 and Susceptible_D5 samples compared to D2. PR-9 (peroxidases) were detected across all sample groups but showed significant increased abundance in Resistant_D5 compared to D2 and Susceptible_D5. PR-10 (hypersensitive-induced reaction proteins) showed increased abundance in both Resistant_D5 and Susceptible_D5 samples relative to D2 (Supplementary Tables 4, 5 and 6).

### Annotation and gene ontology enrichment analysis of barley proteins

Gene ontology (GO) enrichment analysis was used to identify the functional terms associated with the sets of differentially abundant barley proteins, according to their annotations in the reference database for *Hordeum vulgare* MorexV3 (Mascher et al. 2021). The top 20 most significantly enriched protein pathways (FDR < 0.05) for the set of 262 proteins associated with *Ptt* resistance at D5 were linked to translation activities, biosynthesis (peptide, amide, cellular organonitrogen, organic substance), chloroplast, plastids and intracellular organelle (Supplementary Table 7a), the majority of which were associated with chloroplast metabolism (Figure 6). Among the 459 proteins that were associated with *Ptt* susceptibility, the most significantly enriched protein pathways/GO terms were associated with cellular detoxification, chloroplast, plastids, organic acid metabolism, cytoplasm and organonitrogen compound metabolism (Supplementary Table 7b). Regarding the set of 550 proteins that were significantly increased in the Susceptible_D5 samples relative to the D2 samples, the 20 most highly enriched pathways/functional terms were associated with nucleic acid metabolism, amino acid metabolism, organic acid and organic compound metabolism, organonitrogen compound metabolic and cytoplasm (Supplementary Tables 8). For the set of 384 proteins that were significantly increased in the Resistant_D5 samples relative to the D2 samples, the 20 most highly enriched pathways/GO terms were mainly associated with carbohydrate metabolism (Supplementary Tables 9).

**Figure 6.**
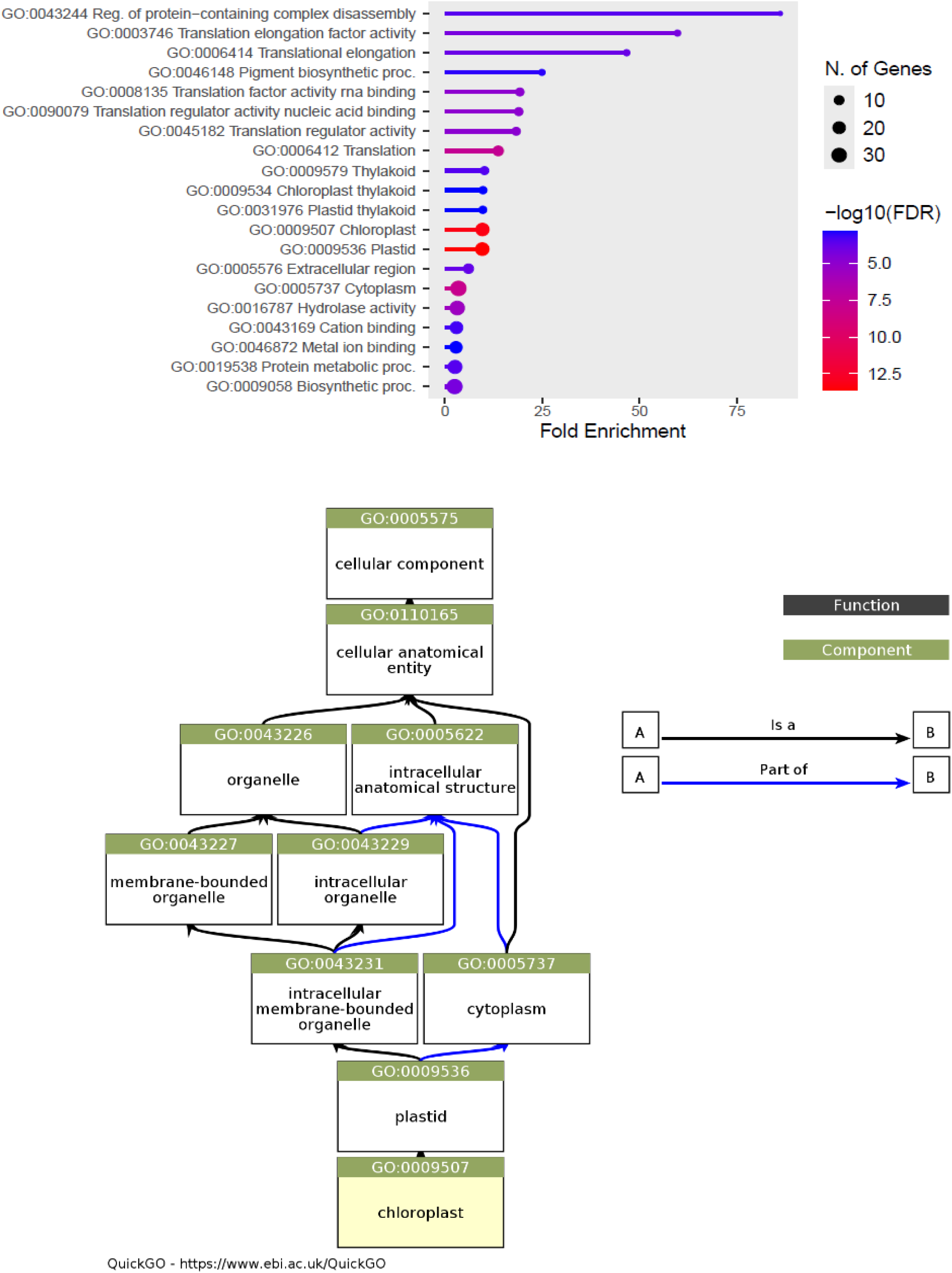
(A) Gene Ontology (GO) enrichment analysis of proteins showed increased abundance in the resistant group compared to the susceptible group of Prior proteomes during *Pyrenophora teres* f. *teres* infection. The bar plot with circles at the end, represents the 20 most significantly enriched GO terms categorized into Biological Processes (BP), Molecular Function (MF), and Cellular Component (CC). The x-axis denotes the GO terms, and the y-axis shows the -log10 (adjusted p-value), indicating the significance of the enrichment. (B) Most of the enriched pathways were associated with chloroplast metabolisms. Colour code represents GO terms categorized into Molecular Function (Function), and Cellular Component (Component).

### The protein profiles of P. teres f. teres during infection

The protein profiles of *Ptt* (*P451, SP23* and *P155*) at D2 and D5 post-inoculation were retrieved from the DIA-MS data. Following quality filtering, a total of 51 *Ptt* protein were identified based on unique *Ptt* peptides (Supplementary Table 1). Of these proteins, 30 were predicted to be effector proteins with potential roles in *Ptt* virulence (Supplementary Table 10). In contrast to the barley proteomic data, several *Ptt* proteins were unique to specific treatment groups, including six proteins that were only identified in all three D5 replicates of the virulent isolates (*P451* and *SP23*) and were absent in the avirulent parent isolates as well as all samples collected at day-2 post-inoculation (Table 2). These proteins were: Avirulence Effector AvrLm4-7 domain-containing protein (E3RDK4), endo-1,4-beta-xylanase (E3S4Z8), protein disulfide-isomerase (E3S8N2), fibronectin type III-like domain-containing protein (E3S0T1), feruloyl esterase C (E3S4G0) and peptidase M14 carboxypeptidase A domain-containing protein (E3RD55). In addition, catalase (E3RV79) and CHRD domain-containing protein (E3S5J2) were only present in all three replicates of virulent and avirulent isolates at 5-days post-inoculation and not detected in the D2 samples.

**Table 2.** List of proteins that were significantly increased in abundance in the *Pyrenophora teres* f. *teres* virulent group (*P451* and *SP23*) compared to the avirulent (*P155*), at five days after inoculation.

| Protein | Gene | Descriptions | *t*-test p-value(-log) | *t*-test q-value | *t*-test Difference |
| --- | --- | --- | --- | --- | --- |
| E3S0T1 | PTT_15710 | Fibronectin type III-like domain-containing protein | 2.447 | 0.018 | 6.797 |
| E3S8N2 | PTT_19353 | Protein disulfide-isomerase | 5.089 | 0.000 | 3.514 |
| E3RP82 | PTT_10429 | Uncharacterized protein | 3.230 | 0.006 | 3.193 |
| E3RT43 | PTT_12150 | Cerato-platanin domain containing protein | 3.936 | 0.002 | 3.114 |
| E3RV79 | PTT_13058 | Catalase | 2.732 | 0.012 | 3.015 |
| E3SAX4 | PTT_20350 | Uncharacterized protein | 3.817 | 0.002 | 2.653 |
| E3S4Z8 | PTT_17674 | Endo-1,4-beta-xylanase | 2.915 | 0.010 | 2.331 |
| E3S766 | PTT_18636 | Uncharacterized protein | 2.489 | 0.018 | 2.199 |
| E3RMG0 | PTT_09632 | Uncharacterized protein | 2.851 | 0.010 | 1.294 |
| E3S5I9 | PTT_17893 | Uncharacterized protein | 1.983 | 0.041 | 0.797 |
| E3RV36 | PTT_13011 | DNA replication factor Cdt1 C-terminal domain-containing protein | 2.078 | 0.036 | -0.354 |
| E3RNR5 | PTT_10228 | Methyltransferase type 11 domain-containing protein | 3.727 | 0.002 | -0.771 |
| E3RPG2 | PTT_10522 | 40S ribosomal protein S23 | 2.144 | 0.033 | -1.327 |

The PCA plot for the *P. teres* proteomics data identified two distinct clusters, in which the barley samples at D2 post-inoculation with virulent *Ptt* (*P451* and *SP23*) clustered separately from the remaining samples, including samples collected D2 post-inoculation with avirulent *Ptt* (*P155*) and all the samples collected at day 5 (Figure 7). Similarly, hierarchical cluster analysis of the protein intensity data and heat map representation of the data (Figure 8) also showed separation of the D2 and D5 samples, with the D2 virulent *Ptt* (*P451* and *SP23*) samples forming a distinct sub-cluster.

**Figure 7.**
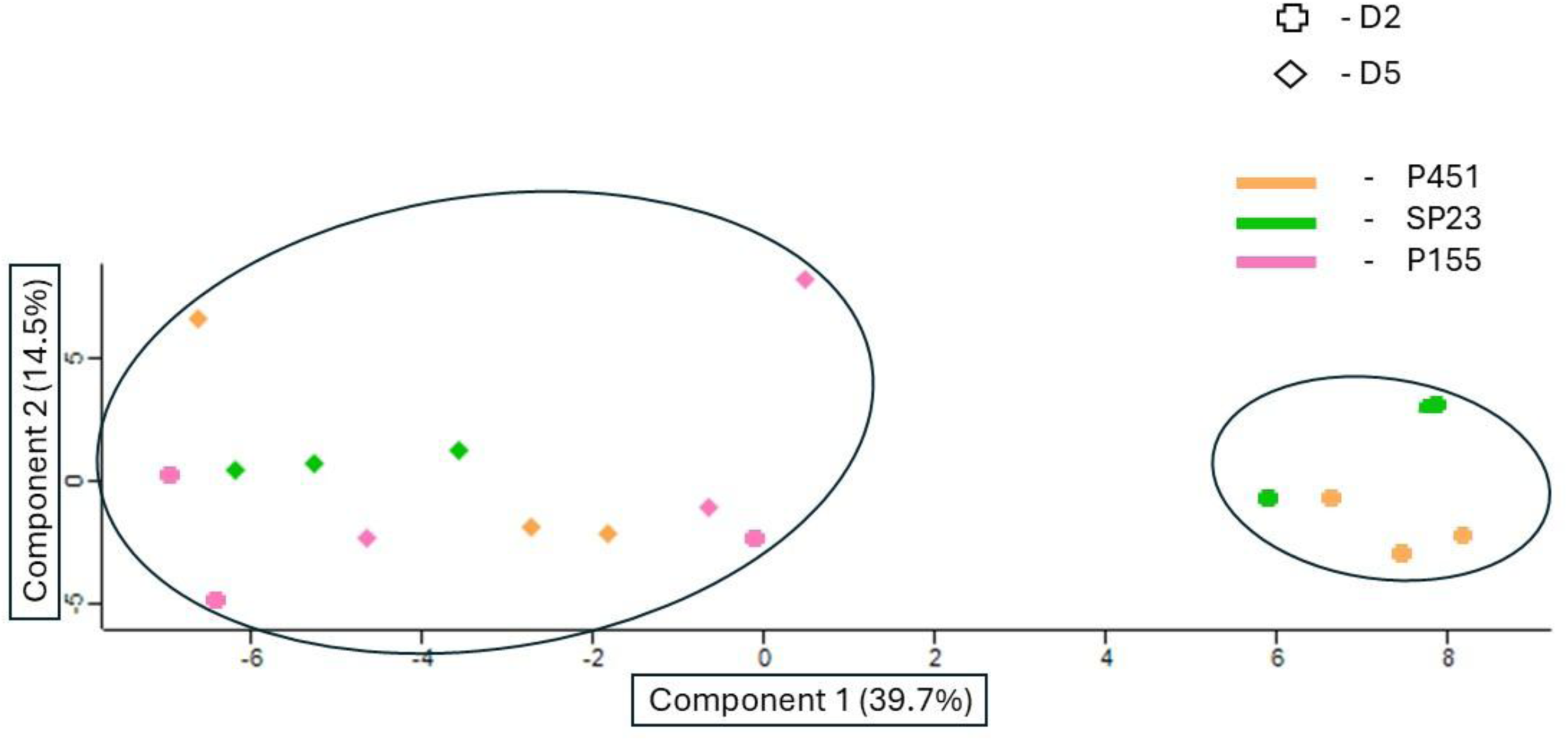
Principal component analysis plot created with 51 *Pyrenophora teres* f. *teres* proteins detected *in planta* during host-pathogen interaction at two and five days after inoculation showing the variations in the proteins of virulent (*P451*) avirulent (*P155*) parental and virulent progeny (*SP23*) isolates. The separation of clusters along PC1 and PC2 highlights the major sources of variation in protein abundance and suggests potential time-point differences between clusters.

**Figure 8.**
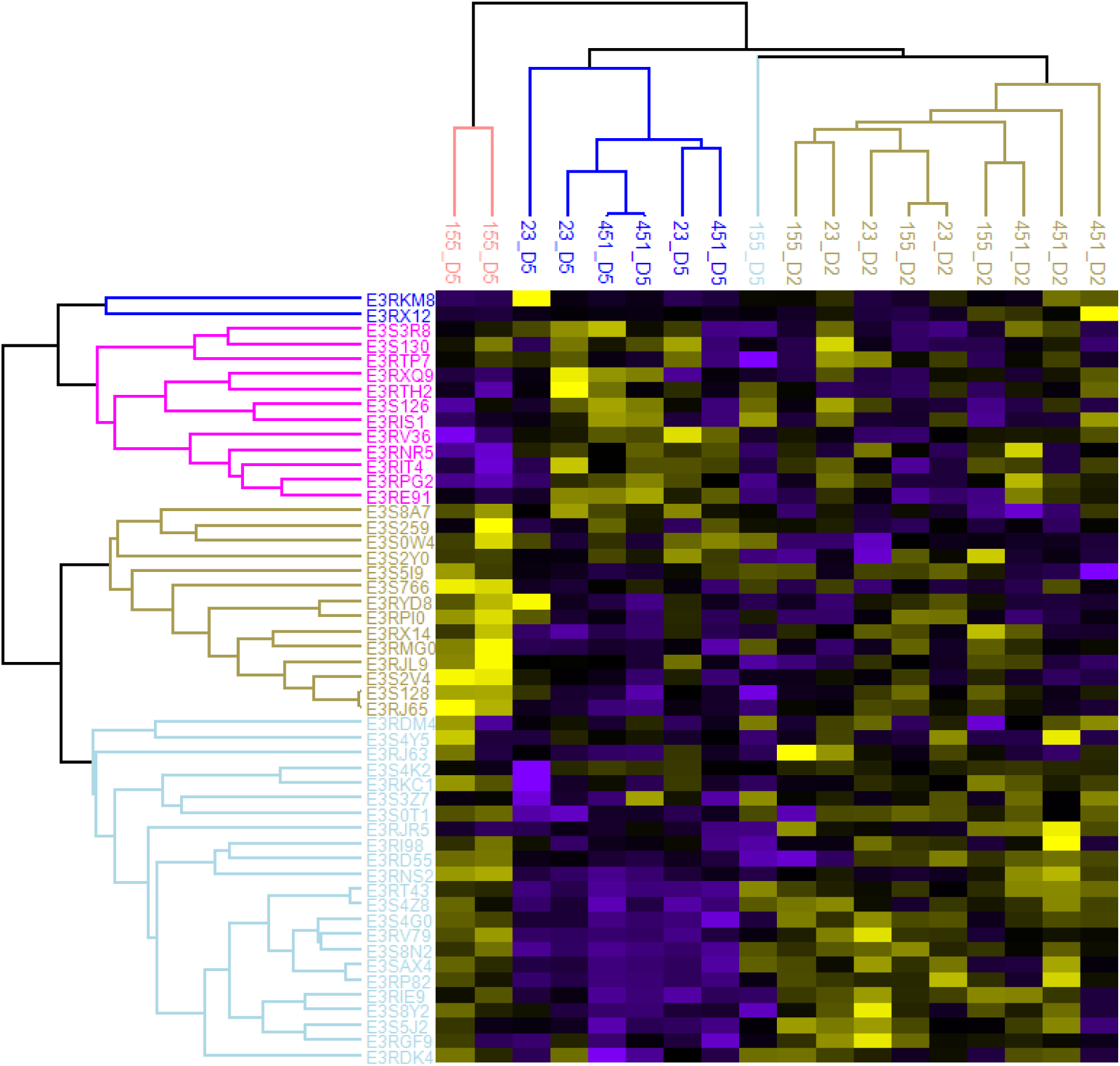
Heatmap displaying the proteins detected in *Pyrenophora teres* f. *teres* (*Ptt*) during host-pathogen interaction at two (D2) and five days (D5) after inoculation in barley cultivar, Prior. Rows represent individual proteins, while columns represent different samples. The colour scale indicates the abundance level, with colour gradient, e.g., yellow to blue representing low to high abundance, respectively. Proteins are grouped into clusters based on their abundance profiles, as determined by hierarchical clustering. Dendrograms along the rows and columns illustrate the hierarchical relationships between proteins and samples, respectively.

Statistical analysis of the *P. teres* proteomics data was then used to identify significantly affected proteins that could be important for the virulence of *P. teres*. Pair-wise comparison of the protein profiles between the virulent (*P451* and *SP23*) and the avirulent (*P155*) samples at D5 post-inoculation identified 13 proteins with significantly (*t*-*tes*t FDR < 0.05) altered abundance (Table 2). Ten proteins were significantly increased in abundance in the virulent group compared to the avirulent group. Pair-wise comparison of the barley samples inoculated with the virulent *Ptt* isolates collected at the D2 and D5 time points identified 11 proteins that were significantly changed in abundance, all of which were more abundant D5 post-inoculation (Table 3).

**Table 3.**
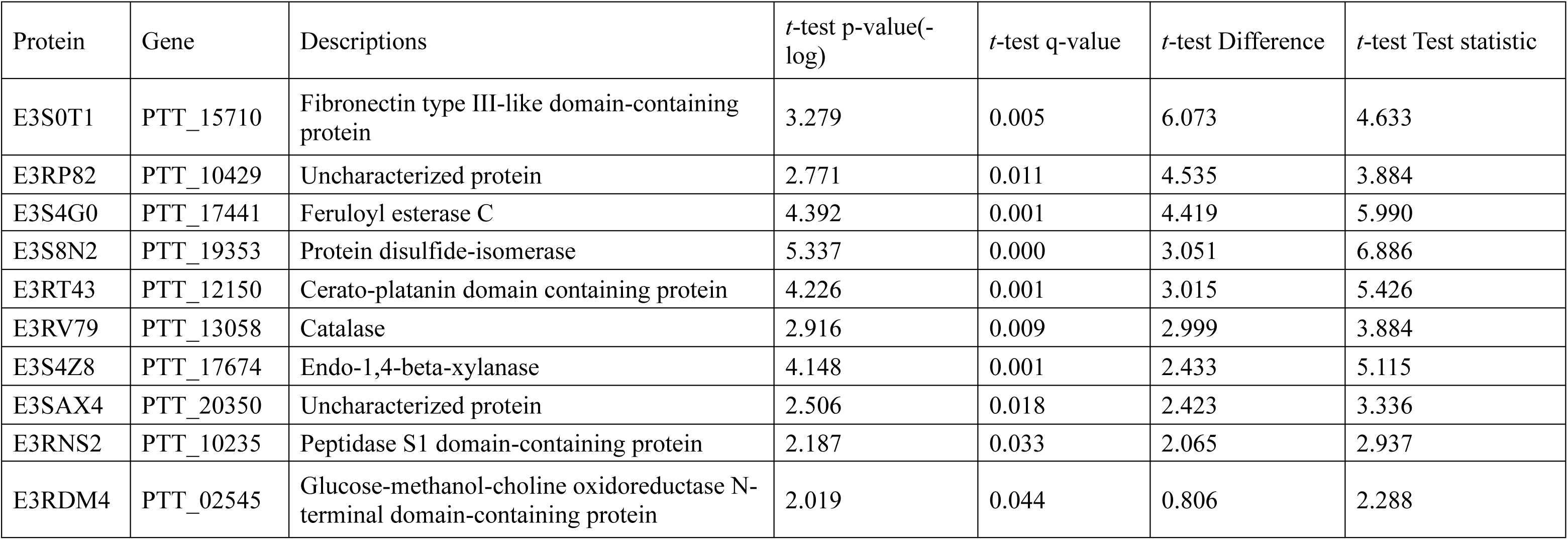
List of proteins that were significantly increased in abundance in the virulent *Pyrenophora teres* f. *teres* group (*P451* and *SP2*3) at five days compared to two days after inoculation.

Proteins within the QTL region identified on *Ptt* chromosome 5 were manually examined. The only protein found in this region was *E3RGF9*, a telomerase reverse transcriptase.

### Annotation and gene ontology enrichment analysis of P. teres f. teres proteins

Gene ontology analysis of *Ptt* proteins that base on Ptt 0-1 (Wyatt et al. 2018) showed increased abundance in the barley samples inoculated with the virulent isolates at D5, compared with the avirulent group, identified a single enriched functional term (FDR < 0.05). Seven of the 10 significant proteins (Table 2) were associated with the term “signal pathways” which had a 7.6-fold enrichment. Regarding comparison between barley samples inoculated with the virulent isolates collected at D5 vs D2, the 11 significant *Ptt* proteins that increased at D5 (Table 3) were classified into six pathways. These significant (FDR < 0.05) pathways were associated with signalling, extracellular regions and catalytic activities and included endo-1,4-beta-xylanase activity, acetylxylan esterase activity, endo-1,4-beta-xylanase activity, and cellulose binding, xylan catabolic process, and cellulase activity.

## Discussion

This study provides an integrated analysis of host and pathogen protein dynamics during the barley-*P. teres* interaction and represents, to our knowledge, the first *in planta* proteomic investigation capturing responses from both organisms simultaneously. By combining pathogen QTL mapping with time-resolved proteomics, we aimed to contextualise a major *Ptt* virulence locus on chromosome 5 within host resistance and susceptibility responses associated with barley variety Prior. This integrated approach enabled the identification of molecular signatures associated with susceptible and resistant interactions while highlighting the complexity of linking genetic loci to downstream protein expression.

Proteomic analysis at D2 and D5 post-inoculation captured distinct phases of the *Ptt* infection process. Early infection responses at D2 were largely similar across all treatments, consistent with shared initial host recognition processes during the brief biotrophic phase. In contrast, pronounced divergence between resistant and susceptible interactions was observed at D5, coinciding with the onset of necrotrophy. Resistant interactions were characterised by increased abundance of proteins associated with translation, amino acid biosynthesis, and chloroplast functions consistent with defence-associated metabolic reprogramming and detoxification (e.g., peroxidases, glutathione-related enzymes, chitinases), whereas susceptible interactions showed enrichment of proteins linked to carbohydrate and organic acid metabolism.

Enrichment analysis revealed that proteins reduced in abundance in the resistant interaction, compared to the susceptible interaction, were predominantly associated with carbohydrate and organic acid metabolism. Increased abundance of host proteins involved in these pathways during susceptible interactions suggests substantial metabolic reprogramming of barley tissue/cells in response to *Ptt* infection. Such metabolic shifts may reflect pathogen-induced modification of host cellular environments, including altered pH regulation and nutrient availability, that favour fungal colonisation and disease development. Enhanced carbohydrate metabolism may indicate reallocation of host carbon resources in susceptible plants, providing readily accessible substrates that support pathogen growth and progression of infection.

Pathogenesis-related (PR) proteins formed a prominent component of the barley response to *Ptt* infection and were differentially abundant between resistant and susceptible interactions. Several PR proteins, including β-1,3-glucanases, peroxidases, chitinases, and germin-like proteins, have been previously implicated in the barley-*P. teres* pathosystem (Hassett et al. 2020; Moolhuijzen et al. 2021). Resistant interactions at D5 were characterised by increased abundance of peroxidases, consistent with effective defence responses, whereas susceptible interactions exhibited higher levels of glucanases, potentially reflecting delayed or less effective activation of defence mechanisms. Chitinases and hypersensitive-induced reaction proteins were elevated in both interaction types at D5, indicating shared defence activation following pathogen recognition, with differences in timing and coordination likely contributing to disease outcome. Minimal PR protein induction at D2 further suggests that strong defence signalling predominantly occurs during later stages of infection.

For this study, we used a bi-parental mapping population to confirm the QTL on chromosome 5 and that the progeny isolates only contain the chromosome 5 QTL. The chromosome 5 QTL spans approximately 209 cM (20 kb based on W1-1 reference genome), representing a large genomic interval that limits fine-scale resolution and confident homology assignment. This region could overlap with previously described *Ptt* or *P. teres* f. *maculata* (*Ptm*) virulence loci based on genomic coordinates and known synteny between the two formae speciales. However, it is not currently possible to determine whether the QTL represents a novel locus or an allelic variant of a previously described region. Detection of this QTL in other barley-*P. teres* studies (Martin et al. 2020; Dahanayaka &Martin 2023) underscores both its potential importance and the need for further fine mapping and comparative genomic analyses.

The parental isolates for this this study were selected based on their contrasting virulences on the barley cultivar Prior and the presence or absence of the major chromosome 5 virulence QTL, while the progeny isolate *SP23* was virulent and had the chromosome 5 QTL. This experimental design enabled comparison of resistant and susceptible host states while minimising background genetic variation in the pathogen. However, despite the large size of the chromosome 5 QTL, only a single gene within the mapped interval, encoding a telomerase reverse transcriptase (E3RGF9), was detected by *in planta* proteomic analysis. The limited recovery of pathogen proteins, relative to host proteins, likely constrained the direct detection of QTL-encoded gene products and reflects a well-recognised challenge in plant-pathogen proteomics. However, this is not unexpected due to the relatively low level of pathogen within a complex host environment (Bindschedler et al. 2009) and the masking effect of more highly abundant host proteins (González-Fernández et al. 2010).

The parental virulent isolate and the single-QTL progeny isolate exhibited broadly similar virulence profiles across the differential set of 24 barley cultivars suggesting that the parental isolate does not carry additional minor-effect QTL, or that, even if such minor QTL are present, the chromosome 5 QTL alone is sufficient to confer the virulence phenotype observed in the parental isolate. Furthermore, any additional minor QTLs present in the parent do not measurably modify disease outcomes under the conditions tested. Collectively, these results support a dominant, largely additive contribution of the chromosome 5 QTL to pathogenicity. Although only a single gene encoding a telomerase reverse transcriptase was detected within the mapped QTL interval by in planta proteomics, this likely reflects the limited detectability of pathogen proteins within host tissue rather than the absence of causal virulence factors. Consistent with this, several virulence-associated proteins identified by proteomics were encoded outside the QTL region, suggesting that the chromosome 5 locus may influence virulence indirectly, for example through regulation of fungal development, growth dynamics, or downstream effector deployment, rather than encoding a detectable effector itself. Nonetheless, 51 specific proteins from *Ptt* were detected, of which six (E3RDK4, E3S4Z8, E3S8N2, E3S0T1, E3S4G0 and E3RD55) were only present in all three replicates at D5 of the virulent isolates and absent in the avirulent isolate. Four of these unique proteins were recognised as apoplastic and one as cytoplasmic effector proteins. Proteins, E3S4Z8, E3S8N2 and E3S0T1 were also significantly more abundant in virulent isolates compared to the avirulent isolate at D5 and the same proteins along with E3S4G0 showed increased abundance on D5 compared to D2 in the virulent isolates. Avirulence effector AvrLm4-7 domain-containing protein (E3RDK4) was reported to play a role in the virulence of hemibiotrophic fungal pathogen, *Leptosphaeria maculans* infection on susceptible *Bassica napus* varieties. AvrLm4-7 suppressed the salicylic acid- and ethylene-dependent signalling pathways coupled with accumulation of reactive oxygen species (ROS) of the host, which are crucial for an effective defence against *L. maculans* (Nováková et al. 2016). Xylanases and FAEs found in *Ptt* are crucial enzymes involved in degradation of cell wall components, such as xylan, hemicellulose and lignin, respectively (Kikot et al. 2009; Balcerzak et al. 2012; Xu et al. 2018).

Protein, E3S8N2; protein disulfide-isomerase (PDI) is well characterised in humans and animals, its importance in fungi remains relatively unknown. A recent study revealed that mutation in a PDI (*FgEps1*) in *Fusarium graminearum* affected growth, asexual and sexual reproduction, stress tolerance and pathogenicity of the pathogen (Liu et al. 2023). Fibronectin type III-like domain-containing protein (E3S0T1) and Peptidase M14 carboxypeptidase A domain-containing protein (E3RD55) are not well studied in plant pathogenic fungi. However, fibronectin type III-like domain-containing proteins present in *Saccharomyces cerevisiae* are reported to be important for the cell-to-cell adhesion (Kraushaar et al. 2015).

In conclusion, this study provides a detailed analysis of the barley-*P. teres* pathosystem, highlighting the dynamic protein changes during early biotrophic and necrotrophic phases of infection. The aim of this study was to identify proteins associated with a single QTL. However, due to the low abundance of fungal proteins, the targeted protein could not be identified. Nevertheless, hundreds of proteins associated with susceptibility and resistance in barley were identified. The identification of proteins including PR proteins involved in resistance and susceptibility, along with the significant enrichment of pathways related to chloroplasts, plastids, and metabolic processes, underscores the complexity of plant-pathogen interactions. Most of the fungal proteins identified are potential effector proteins hence, these proteins would provide valuable information on the pathogen’s ability to manipulate host cellular processes, suppress immune responses, and facilitate infection.

## Supporting information

Supplementary_QTL mapping

## Funding

QTL mapping segment of this study was funded by the Australian Research Council Linkage grant ARC_LP220100084.

## Competing Interests

The authors have no relevant financial or non-financial interests to disclose.

## Author Contributions

B.D. and A.M. performed the QTL analysis. B.D. and S.B. performed the proteomic experiments. R.W. performed the proteomic sample analyses. B.D. and R.W. performed proteomic data analyses. B.D. prepared and revised the manuscript, B.D. and R.W. revised the manuscript after reviewers’ suggestions. S.B., R.W., J.H. and A.M. reviewed the manuscript.

## Data Availability

The mass spectrometry proteomics data have been deposited to the ProteomeXchange Consortium via the PRIDE partner repository with the dataset identifier PXD066431.

## Ethics, Consent to Participate, and Consent to Publish declarations

Not applicable

## Identification and Voucher Specimens for plant materials

None

## Notes

### Competing Interest Statement

The authors have declared no competing interest.

## Reference

Anderson JP, Hane JK, Stoll T, et al. (2016). Proteomic analysis of Rhizoctonia solani identifies infection-specific, redox associated proteins and insight into adaptation to different plant hosts. Molecular & Cellular Proteomics 15: 1188–1203

Bakonyi J, Seress D, Nagy ZÁ, et al. (2024). Virulence Spectra of Hungarian Pyrenophora teres f. teres Isolates Collected from Experimental Fields Show Continuous Variation without Specific Isolate× Barley Differential Interactions. Journal of Fungi 10: 184

Balcerzak M, Harris LJ, Subramaniam R, et al. (2012). The feruloyl esterase gene family of Fusarium graminearum is differentially regulated by aromatic compounds and hosts. Fungal biology 116: 478–488

Balotf S, Wilson R, Tegg RS, et al. (2021). In planta transcriptome and proteome profiles of Spongospora subterranea in resistant and susceptible host environments illuminates regulatory principles underlying host–pathogen interaction. Biology 10: 840

Bindschedler LV, Burgis TA, Mills DJ, et al. (2009). In planta proteomics and proteogenomics of the biotrophic barley fungal pathogen Blumeria graminis f. sp. hordei. Molecular & Cellular Proteomics 8: 2368–2381

Böhmer M, Colby T, Böhmer C, et al. (2007). Proteomic analysis of dimorphic transition in the phytopathogenic fungus Ustilago maydis. Proteomics 7: 675–685

Casey T, Solomon PS, Bringans S, et al. (2010). Quantitative proteomic analysis of G-protein signalling in Stagonospora nodorum using isobaric tags for relative and absolute quantification. Proteomics 10: 38–47

Churchill GA, Doerge RW (1994). Empirical threshold values for quantitative trait mapping. Genetics 138: 963–971

Clare SJ, Wyatt NA, Brueggeman RS, et al. (2020). Research advances in the Pyrenophora teres–barley interaction. Molecular Plant Pathology 21: 272–288

Dahanayaka, Vaghefi N, Snyman L, et al. (2021a). Investigating in vitro mating preference between or within the two forms of Pyrenophora teres and its hybrids. Phytopathology® 111: 2278–2286

Dahanayaka B, Martin A (2023). Multi-parental fungal mapping population study to detect genomic regions associated with Pyrenophora teres f. teres virulence. Scientific Reports 13: 9804

Dahanayaka BA, Snyman L, Vaghefi N, et al. (2022a). Using a hybrid mapping population to identify genomic regions of Pyrenophora teres associated with virulence. Frontiers in Plant Science 13: 925107

Dahanayaka BA, Snyman L, Vaghefi N, et al. (2022b). Using a hybrid mapping population to identify genomic regions of *Pyrenophora teres* associated with virulence. Front Plant Sci: 2059

Dahanayaka BA, Vaghefi N, Knight N, et al. (2021b). Population structure of *Pyrenophora teres* f. *teres* barley pathogens from different continents. Phytopathology 111: 2118–2129

De Wit PJ, Laugé R, Honée G, et al. (1997). Molecular and biochemical basis of the interaction between tomato and its fungal pathogen Cladosporium fulvum. Antonie Van Leeuwenhoek 71: 137–141

De Wit PJ, Mehrabi R, Van den Burg HA, et al. (2009). Fungal effector proteins: past, present and future. Molecular plant pathology 10: 735–747

Doerge RW, Churchill G (1996). Permutation tests for multiple loci affecting a quantitative character. Genetics 142: 285–294

Ellwood SR, Liu Z, Syme RA, et al. (2010). A first genome assembly of the barley fungal pathogen Pyrenophora teres f. teres. Genome biology 11: R109

Ellwood SR, Piscetek V, Mair WJ, et al. (2019). Genetic variation of *Pyrenophora teres* f. *teres* isolates in Western Australia and emergence of a Cyp51A fungicide resistance mutation. Plant Pathology 68: 135–142

Ellwood SR, Syme RA, Moffat CS, et al. (2012). Evolution of three Pyrenophora cereal pathogens: recent divergence, speciation and evolution of non-coding DNA. Fungal Genetics and Biology 49: 825–829

Fowler RA, Platz GJ, Bell KL, et al. (2017). Pathogenic variation of Pyrenophora teres f. teres in Australia. Australasian Plant Pathology 46: 115–128

Friesen TL, Faris JD, Solomon PS, et al. (2008). Host-specific toxins: effectors of necrotrophic pathogenicity. Cellular Microbiology 10: 1421–1428

Friesen TL, Meinhardt SW, Faris JD (2007). The Stagonospora nodorum-wheat pathosystem involves multiple proteinaceous host-selective toxins and corresponding host sensitivity genes that interact in an inverse gene-for-gene manner. The Plant Journal 51: 681–692

Ge SX, Jung D, Yao R (2020). ShinyGO: a graphical gene-set enrichment tool for animals and plants. Bioinformatics 36: 2628–2629

González-Fernández R, Aloria K, Valero-Galván J, et al. (2014). Proteomic analysis of mycelium and secretome of different Botrytis cinerea wild-type strains. Journal of proteomics 97: 195–221

González-Fernández R, Prats E, Jorrín-Novo JV (2010). Proteomics of plant pathogenic fungi. BioMed Research International 2010: 932527

Hassett K, Ellwood SR, Zulak KG, et al. (2020). Analysis of apoplastic proteins expressed during net form net blotch of barley. Journal of Plant Diseases and Protection 127: 683–694

Hebert T (1971). The perfect stage of Pyricularia grisea.

Hughes CS, Moggridge S, Müller T, et al. (2019). Single-pot, solid-phase-enhanced sample preparation for proteomics experiments. Nature protocols 14: 68–85

Jacques S, Lenzo L, Stevens K, et al. (2021). An optimized sporulation method for the wheat fungal pathogen Pyrenophora tritici-repentis. Plant Methods 17: 52

Jones JD, Dangl JL (2006). The plant immune system. Nature 444: 323–329

Kikot GE, Hours RA, Alconada TM (2009). Contribution of cell wall degrading enzymes to pathogenesis of Fusarium graminearum: a review. Journal of basic microbiology 49: 231–241

Kraushaar T, Brückner S, Veelders M, et al. (2015). Interactions by the fungal Flo11 adhesin depend on a fibronectin type III-like adhesin domain girdled by aromatic bands. Structure 23: 1005–1017

Langmead B, Salzberg SL (2012). Fast gapped-read alignment with Bowtie 2. Nature methods 9: 357–359

Lehmensiek A, Bovill W, Wenzl P, et al. (2009) Genetic mapping in the Triticeae. In: Genetics and Genomics of the Triticeae. Springer, pp 201–235

Leiva-Mora M, Capdesuñer Y, Villalobos-Olivera A, et al. (2024). Uncovering the Mechanisms: The Role of Biotrophic Fungi in Activating or Suppressing Plant Defense Responses. Journal of Fungi 10: 635

Li J, Zhang X, Li L, et al. (2018). Proteomics analysis of SsNsd1-mediated compound appressoria formation in Sclerotinia sclerotiorum. International Journal of Molecular Sciences 19: 2946

Lightfoot DJ, Able AJ (2010). Growth of Pyrenophora teres in planta during barley net blotch disease. Australasian Plant Pathology 39: 499–507

Liu K, Wang X, Li Y, et al. (2023). Protein Disulfide Isomerase FgEps1 Is a Secreted Virulence Factor in Fusarium graminearum. Journal of Fungi 9: 1009

Manly KF, Cudmore J, Robert H, Meer JM (2001). Map Manager QTX, cross-platform software for genetic mapping. Mammalian Genome 12: 930–932

Marra R, Li H, Barbetti M, et al. (2010). Proteomic analysis of the interaction between Brassica napus cv. Surpass 400 and virulent or avirulent isolates of Leptosphaeria maculans. Journal of Plant Pathology: 89–101

Martin A, Moolhuijzen P, Tao Y, et al. (2020). Genomic regions associated with virulence in Pyrenophora teres f. teres identified by genome-wide association analysis and biparental mapping. Phytopathology 110: 881–891

Mascher M, Wicker T, Jenkins J, et al. (2021). Long-read sequence assembly: a technical evaluation in barley. The Plant Cell 33: 1888–1906

Mayer K, Waugh R, Langridge P, et al. (2012). A physical, genetic and functional sequence assembly of the barley genome. Nature 491: 711–716

Meng Q, Gupta R, Min CW, et al. (2019). Proteomics of Rice—Magnaporthe oryzae interaction: what have we learned so far? Frontiers in Plant Science 10: 1383

Moolhuijzen P, Lawrence JA, Ellwood SR (2021). Potentiators of disease during barley infection by Pyrenophora teres f. teres in a susceptible interaction. Molecular plant-microbe interactions 34: 779–792

Murray G, Brennan J (2010). Estimating disease losses to the Australian barley industry. Australasian Plant Pathology 39: 85–96

Nováková M, Šašek V, Trdá L, et al. (2016). L eptosphaeria maculans effector AvrLm4-7 affects salicylic acid (SA) and ethylene (ET) signalling and hydrogen peroxide (H2O2) accumulation in B rassica napus. Molecular Plant Pathology 17: 818–831

Paper JM, Scott-Craig JS, Adhikari ND, et al. (2007). Comparative proteomics of extracellular proteins in vitro and in planta from the pathogenic fungus Fusarium graminearum. Proteomics 7: 3171–3183

Rauwane ME, Ogugua UV, Kalu CM, et al. (2020). Pathogenicity and virulence factors of Fusarium graminearum including factors discovered using next generation sequencing technologies and proteomics. Microorganisms 8: 305

Smedegard-Petersen V (1976). Pathogenesis and genetics of net-spot blotch and leaf stripe of barley caused by Pyrenophora teres and Pyrenophora graminea.

Sperschneider J, Dodds PN (2022). EffectorP 3.0: prediction of apoplastic and cytoplasmic effectors in fungi and oomycetes. Molecular plant-microbe interactions 35: 146–156

Thynne E, Ali H, Seong K, et al. (2024). An array of Zymoseptoria tritici effectors suppress plant immune responses. Molecular Plant Pathology 25: e13500

Tian J, Chen C, Sun H, et al. (2021). Proteomic analysis reveals the importance of exudates on sclerotial development in Sclerotinia sclerotiorum. Journal of Agricultural and Food Chemistry 69: 1430–1440

Tyanova S, Temu T, Sinitcyn P, et al. (2016). The Perseus computational platform for comprehensive analysis of (prote) omics data. Nature methods 13: 731–740

Van Os H, Stam P, Visser RG, et al. (2005). RECORD: a novel method for ordering loci on a genetic linkage map. Theoretical and Applied Genetics 112: 30–40

Wang J, Diaz J, Hua K, et al. (2024). Proteomic identification of apoplastic proteins from rice, wheat, and barley after Magnaporthe oryzae infection. Phytopathology Research 6: 55

Wang S (2012). Windows QTL cartographer 2.5. (No Title):

Weiland P, Altegoer F (2021). Identification and characterization of two transmembrane proteins required for virulence of Ustilago maydis. Frontiers in Plant Science 12: 669835

Wyatt NA, Richards JK, Brueggeman RS, et al. (2018). Reference assembly and annotation of the Pyrenophora teres f. teres isolate 0-1. G3: Genes, Genomes, Genetics 8: 1–8

Xu M, Gao X, Chen J, et al. (2018). The feruloyl esterase genes are required for full pathogenicity of the apple tree canker pathogen Valsa mali. Molecular Plant Pathology 19: 1353–1363

